# The effects of demography and genetics on the neutral distribution of quantitative traits

**DOI:** 10.1101/421008

**Authors:** Evan M. Koch

## Abstract

Neutral models for quantitative trait evolution are useful for identifying phenotypes under selection in natural populations. Models of quantitative traits often assume phenotypes are normally distributed. This assumption may be violated when a trait is affected by relatively few genetic variants or when the effects of those variants arise from skewed or heavy-tailed distributions. Traits such as gene expression levels and other molecular phenotypes may have these properties. To accommodate deviations from normality, models making fewer assumptions about the underlying trait genetics and patterns of genetic variation are needed. Here, we develop a general neutral model for quantitative trait variation using a coalescent approach by extending the framework developed by Schraiber and Landis (2015). This model allows interpretation of trait distributions in terms of familiar population genetic parameters because it is based on the coalescent. We show how the normal distribution resulting from the infinitesimal limit, where the number of loci grows large as the effect size per mutation becomes small, depends only on expected pairwise coalescent times. We then demonstrate how deviations from normality depend on demography through the distribution of coalescence times as well as through genetic parameters. In particular, population growth events exacerbate deviations while bottlenecks reduce them. This model also has practical applications, which we demonstrate by designing an approach to simulate from the null distribution of Q_ST_, the ratio of the trait variance between subpopulations to that in the overall population. We further show that it is likely impossible to distinguish sparsity from skewed or heavy-tailed distributions of mutational effects using only trait values sampled from a population. The model analyzed here greatly expands the parameter space for which neutral trait models can be designed.

## 2 Introduction

Neutral models of quantitative traits provide a null distribution against which various goodness-of-fit tests can be used to test for the action of natural selection (Lande, 1976; Leinonen *et al.*, 2013). Neutral models can also clarify the effects of purely neutral forces such as genetic drift and mutation on trait distributions (Lynch and Hill, 1986). A common approach is to first model phenotypes as normally distributed, either among offspring within a family, among members of a population, or among species (Turelli, 2017). Indeed, it has been suggested that the normality assumption is the defining characteristic of quantitative genetics (Rice, 2004). This might be justified if phenotypes are influenced by a large number of sufficiently independent Mendelian factors (Fisher, 1918), or normality may simply appear approximately true in practice.

Neutral models for quantitative traits have been developed in a variety of contexts. The goal of these models is to ask to whether phenotypic differentiation between groups can be reasonably explained by processes other than natural selection. On macroevolutionary time scales, models stemming from Lande (1976) have used Brownian motion to describe the evolution of the mean value of a quantitative trait in a population. These models are used in statistical methods to test for extreme trait divergence between species (Turelli *et al.*, 1988), test for phylogenetic signal in trait distributions (Freckleton *et al.*, 2002), and correct for phylogenetic dependence when calculating correlations between traits (Felsenstein, 1985). On shorter time scales, neutral distributions assuming multivariate normality of trait values also underlie tests for spatially varying selection in structured populations such as the method developed by Ovaskainen *et al.* (2011). Other neutral models for quantitative traits have not assumed normality (Chakraborty and Nei, 1982; Lynch and Hill, 1986; Lande, 1992), and the dynamics of phenotypic evolution are examined forwards in time as a balance between mutation creating variance, migration spreading variance among subpopulations, and fixation removing it. However, these studies were limited to simple models of population structure and history. Backwards-in-time, coalescent models would allow for more general demographic scenarios.

Under the normality assumption, quantitative trait dynamics can be modeled without concern for the number of causal loci influencing the trait, the genealogies at these sites, or the distribution of mutational effects (the distribution that new mutations affecting the trait draw their effects from). However, heritable phenotypic variation is ultimately due to discrete mutations at discrete locations in the genome, and how the phenotypic variance arising from these mutations is distributed depends on the genealogies at these loci. For instance, the distribution of genealogies in the genome might be strongly influenced by recent population growth and the distribution of mutational effects could be skewed for biological reasons inherent in the details of a particular developmental pathway. When the number of mutations affecting a trait is large, the central limit theorem ensures that the distributions of genealogies and mutational effects can be ignored, but a full model of phenotypic variation would have to include them. Importantly, deviations from normality may affect the outcomes of goodness-of-fit tests that necessarily aim to identify outliers from a normal model.

A more inclusive model of neutral phenotypic variation can begin by considering the genealogies at causal loci. The principle modeling framework for genealogical variation is the coalescent process (Wakeley, 2008), but few studies have connected the coalescent to quantitative genetics. Whitlock (1999) used coalescent theory to argue that measures of phenotypic (*Q*_*ST*_) and genetic (*F*_*ST*_) differentiation have the same expected value given general models of population subdivision. By simulating from the coalescent with recombination, Griswold *et al.* (2007) investigated the effects of shared ancestry and linkage disequilibrium on the genetic covariance matrix for a set of traits (**G** matrix). They found that linkage disequilibrium and small numbers of causal loci can cause phenotypic covariances not predicted by the mutational covariance matrix. Mendes *et al.* (2018) also used a quantitative trait model based on the coalescent to show that using a species tree based on population split times can lead to, among other problems, increased false positive rates in phylogenetic comparative methods. Although not explicitly connected to the coalescent, Ovaskainen *et al.* (2011) developed their test for spatially varying selection by assuming that the covariance in trait values, conditional on the G-matrix in the ancestral population, depends only on the pairwise coancestry coefficients, which have a clear interpretation in terms of the coalescent process (Slatkin, 1991).

Two studies have asked how the shape of the distribution of mutational effect sizes, beyond just the mutational variance, impacts trait distributions. Khaitovich *et al.* (2005) modeled the evolution of gene expression values on phylogenetic trees assuming a single non-recombining causal locus but allowed for an arbitrary distribution of mutational effects. Using this model they were able to detect deviations from normality consistent with asymmetries in the distribution of mutational effects on gene expression in great apes. More recently, Schraiber and Landis (2015) developed a similar general model of quantitative trait evolution at the population level based on the coalescent and allowing for any number of causal loci. They derived the characteristic function for the distribution of phenotypic values in a sample and showed how such values can deviate strongly from normality when the number of loci is small or the mutational distribution has heavy tails. Schraiber and Landis (2015) note that the possibility for multimodal trait distributions could lead to incorrect inferences of divergent selection within populations.

Schraiber and Landis (2015) derived their results for a panmictic, constant-size population. Natural populations rarely have stable population sizes and show considerable spatial structure, and it is unclear how these violations of the constant-size, panmictic model might influence deviations from normality. We take advantage of the ability of coalescent theory to handle nonequilibrium demographies and population structure to relax these modeling assumptions.

Extending coalescent models of quantitative traits to structured populations is important because the analysis of structured populations provides an opportunity to infer the incidence of local adaptation or stabilizing selection. Due to the build up of linkage disequilibrium between alleles affecting a trait, the neutral divergence in trait values among different subpopulations in a structured population is about as variable as the variance in allele frequencies (Rogers and Harpending, 1983). The *Q*_*ST*_ /*F*_*ST*_ paradigm was developed to test whether trait divergence in structured populations could be explained by neutral forces alone (Spitze, 1993; Whitlock, 2008; Leinonen *et al.*, 2013). *Q*_*ST*_, defined as the ratio of the trait variance between subpopulations to the total trait variance, is compared to *F*_*ST*_, which measures the same property for genetic variation and is calculated using neutral markers to provide a null distribution. If the observed *Q*_*ST*_ is sufficiently far from the null expectation, it is concluded that natural selection has acted. Ovaskainen *et al.* (2011) developed a modern extension of the *Q*_*ST*_ /*F*_*ST*_ paradigm for genetic values measured in breeding experiments and Berg and Coop (2014) also extended the paradigm to make use of genetic values computed from GWAS summary statistics. An advantage of the Berg and Coop (2014) approach is that by using computing genetic values from GWAS loci it makes no assumptions about normality at the population level. However, since suitably sized GWAS’s have only been performed in humans, the approach has not yet been extended to other species. Understanding the neutral distribution of trait values is therefore necessary for the development of goodness-of-fit tests that are applicable to populations with complicated histories and traits with sparse genetic architectures.

We generalize the work of Schraiber and Landis (2015) by deriving the form of the moment generating function (mgf) for arbitrary distributions of coalescent times (e.g. those arising under exponential growth or an island model of migration). The key result of Schraiber and Landis (2015), the characteristic function of the sampling distribution of phenotypic values, is a special case of this general generating function. We then show how a normal model arises by taking the infinitesimal limit where the effect size per mutation becomes small as the number of loci potentially affecting the trait becomes large. We then calculate the third and fourth central moments of the trait distribution in panmictic populations to illustrate how departures from normality depend both on genetic parameters and genealogical distributions. For instance, in exchangeable populations the expected third central moment is proportional to the third noncentral moment of the mutational distribution times the expected time to the first coalescent event in a sample of size three.

Finally, we discuss the consequences of these results for *Q*_*ST*_ tests and the inference of genetic parameters. We find an improved null distribution that can be derived simply by using the normal distribution that arises in the infinitesimal limit of our coalescent model. Additionally, we show that it is likely not possible to infer most features of the mutational distribution using only trait values sampled from a population. Future work will be necessary to develop tests for selection that take into account both demography and genetic parameters, but the model developed here provides the groundwork for such an undertaking.

## 3 Model

In the model we investigate here, there are *L* unlinked potentially causal loci at which mutations influence the trait value. Following Kimura (1969), an infinite number of mutations are possible within each locus and the rate per unit of coalescent time at which mutations affecting the trait (causal mutations) arise is 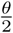. That is, 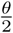 is the mutation rate for the entire locus and not per nucleotide. An approximation for when at most one mutation per locus is likely is considered in Section 6. Mutations are randomly assigned effects from a distribution of effect sizes, and effects are additive both within and between loci. The moment generating function (mgf) of this distribution is written as *ψ* and the *i*^*th*^ non-central moment is *m*_*i*_. Individuals are haploid and the sum of all mutations occurring in an individual’s history determines the trait value of the individual. An extension to diploidly would be straightforward but is not considered here. Correlations between individuals arise when mutations fall on shared portions of genealogies at specific loci. Because the loci are unlinked we assume their genealogies are independent. This model is shown schematically in Figure 1.

**Figure 1:**
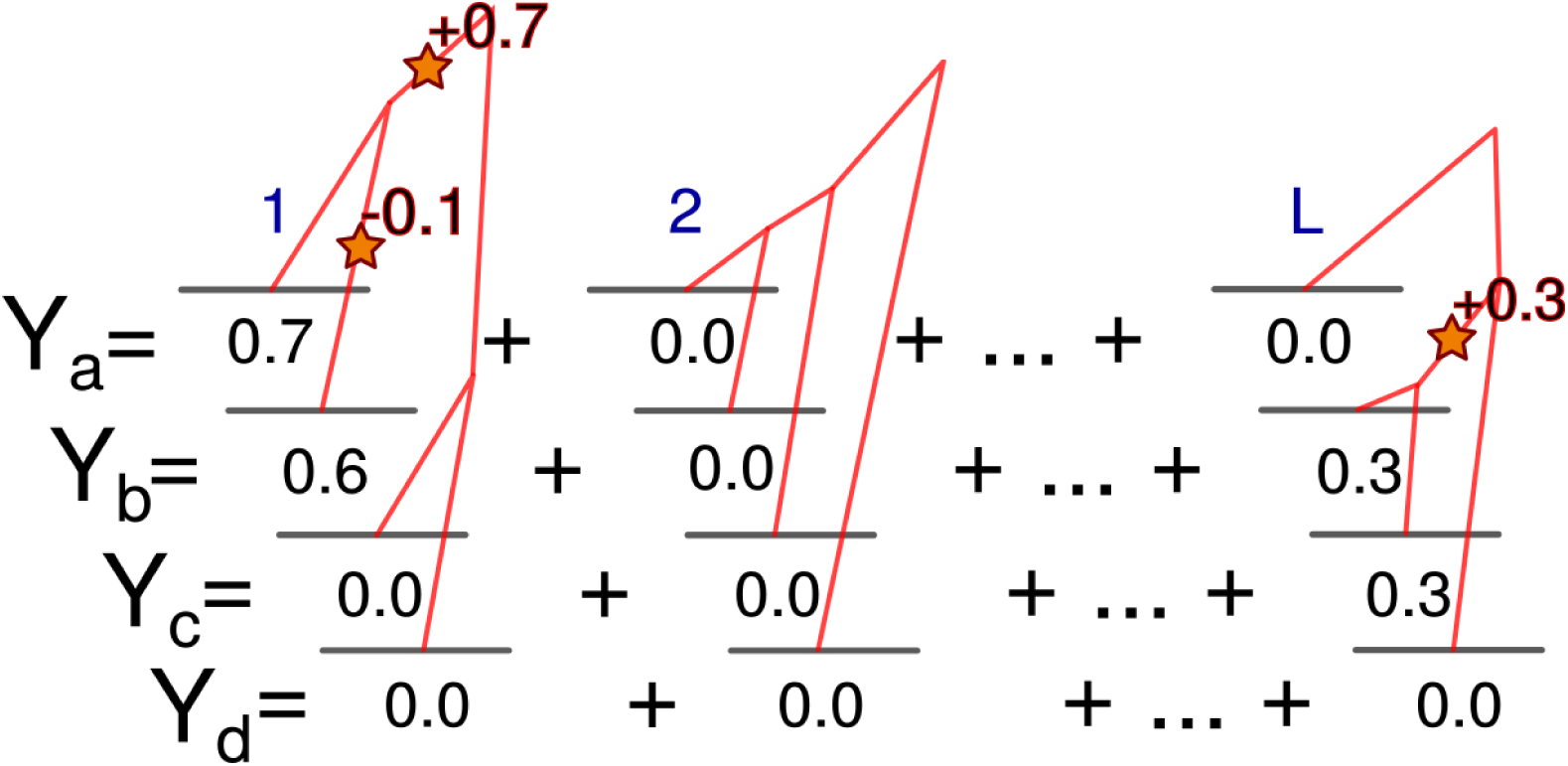
A schematic representation of the model for how trait distributions arise from genealogical and mutational processes. *L* loci potentially affect the trait in a set of individuals and have independent genealogies. Mutations occur within loci as a Poisson process and act additively to give individual trait values. Many loci with the potential to affect the trait may receive no mutations.

The genealogy at a locus is represented by the random vector of branch lengths, **T**. An element *T*_*ω*_ of **T** is the branch length subtending only individuals in the set *ω* and no others in the sample. For example, *T*{*_a,b_*} is the length of the branch subtending only individuals *a* and *b*. If a branch subtending only *a* and *b* does not exist for a given genealogy, *T*{*_a,b_*} is set to zero. In this way **T** encodes both the branch lengths and the topology of a genealogy. Ω is the set of all possible branches. If there are three sampled individuals, *a, b*, and *c*, then Ω = {{*a*}, {*b*}, {*c*}, {*a, b*}, {*a, c*}, {*b, c*}} and **T** = (*T*{*_a_*}, *T*{*_b_*}, *T*{*_c_*}, *T*{*_a,b_*}, *T*{*_a,c_*}, *T*{*_b,c_*}). The mgf for the distribution of branch lengths is denoted *φ***T**.

Phenotypic trait values are the random quantities we are interested in and result from mutations occuring along the branches of genealogies. We will hereafter refer to the phenotypic trait simply as trait values and ignore any environmental component. The random vector of trait values in the sampled individuals is **Y**. If we had sampled individuals *a, b*, and *c*, then **Y** = (*Y*_*a*_, *Y*_*b*_, *Y*_*c*_). The contribution to the trait values from a single locus is the change relative to the value in the most recent common ancestor (MRCA) of the sample at that locus. Since we do not know the ancestral value, we cannot directly observe the change in trait values. Thus, for a trait controlled by multiple loci, **Y** is the sum over contributions from these loci, each measured with respect to an arbitrary value. However, **Y** is sufficient to determine measurable quantities such as differences in trait values between individuals as well as the sample variance. The moment generating functions for the distribution of trait values is denoted *φ***Y**.

Here we refer to the genetic parameters of a trait as the combination of quantities not influenced by the genealogical process: (*L*, 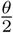, *ψ*). Another quantity useful for describing a trait’s distribution is its sparsity. Sparsity should reflect how many mutations segregating in the population influence the trait, with a more ‘sparse’ trait being one affected by fewer segregating mutations. Formally, we measure sparsity as the average number of pairwise differences between two randomly chosen haplotypes at loci affecting the trait. A trait with fewer causal pairwise differences is more sparse. Sparsity thus depends both on the genetic parameters through the mutation rate, the number of potentially causal loci, and the distribution of coalescence times.

In populations of exchangeable individuals, a useful way to summarize the distribution of genealogies is through the moments of 𝕋 *_k,n_* which denotes the amount of time that *k* lineages remain in the genealogy of a sample of size *n*. The pairwise coalescent time between a lineage in individual *i* and in individual *j* is written as 𝒯 *_i,j_*. When considering structured populations, 𝒯 *_a,b_* is also used to denote the coalescence time between a randomly chosen lineage from subpopulation *a* and a randomly chosen lineage from subpopulation *b*. A final set of quantities are defined for sums of branch lengths. Let *τ*_*a*_+*_b_* be the sum of all branches ancestral to both *a* and *b*, and *τ*_*a/b*_ be the sum of all branches ancestral to *a* but not *b*. Extensions of this for more than two individuals are also used. The same notation is used when referring to sets of branch indices. So Ω_*a*+*b*_ and Ω*_a/b_* would be the sets of branches added to get *τ*_*a*+*b*_ and *τ*_*a/b*_ respectively.

## 4 The moment generating function for the distribution of trait values

We first derive the mgf of the distribution of trait values following closely the approach of Schraiber and Landis (2015) and Khaitovich *et al.* (2005), but generalizing to arbitrary demographies and population structure. We consider the distribution of trait values over evolutionary realizations of the combined random processes of drift and mutation. The probability distribution for a trait is complex in its general form. There is a point mass at zero corresponding the possibility that no mutations affecting the trait occur, and mutational effects could be drawn from discrete or continuous distributions. Correlations between individuals arise because of shared history in the genealogies at individual loci. An analytical expression for the probability distribution of trait values does not exist except in certain limits such as when the number of mutations affecting the trait becomes large.

However, even in the absence of a probability distribution function we can use the mgf approach to learn something about the distribution of trait values. Following the definition of the mgf for a vector-valued random variable, the mgf for a trait controlled by a single nonrecombining locus is

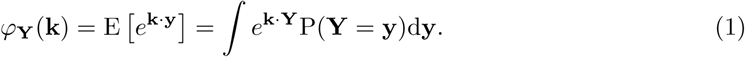

The vector **k** contains dummy variables for each individual, and the whole operation is an intergral transform of the probability distribution of trait values. Equation (1) can be rewritten by conditioning on the genealogy to give

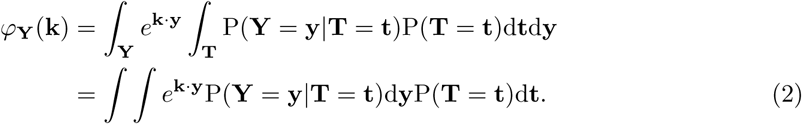

To proceed it is necessary to make assumptions about the mutational process. The first is that mutations occur as a Poisson process along branches and the second is that mutations at a locus are additive. Under these assumptions, the changes in the trait value along each branch are conditionally independent given the branch lengths. Khaitovich *et al.* (2005) and Schraiber and Landis (2015) noted that this describes a compound Poisson process. The mgf of a compound Poisson process with rate *λ* over time *t* is exp(*λt*(*ψ*(*k*) *-*1)), where *ψ* is the mgf of the distribution of the jump sizes caused by events in the Poisson process. In this case the jump sizes are the effects on the trait value caused by new mutations. Using this expression for the mgf of a compound Poisson process, along with the fact that the mgf of two perfectly correlated random variables with the same marginal distribution is *φ*_*X*1_ (*k*_1_ + *k*_2_), we can rewrite equation (2) as

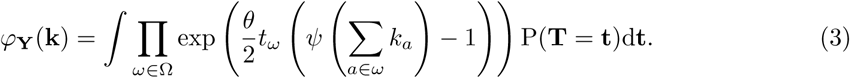

We can recognize equation (3) as the mgf of **T** with the dummy variable *s*_*ω*_ for branch *T*_*ω*_ set to 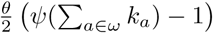. Or,

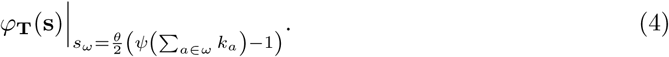

Thus, if the mgf of the distribution of branch lengths is known, equation (3) allows us to obtain the mgf of the trait values through a simple substitution. When the trait is controlled by *L* independent loci, the full mgf, *φ***Y**(**k**), is obtained by raising equation (4) to the power *L*. This result obviates the need for separate derivations for particular models of population history and structure. Lohse *et al.* (2011) derived the mgf of the genealogy in various population models including migration and splitting of subpopulations. Using their result for a single population we could obtain equation (1) of Schraiber and Landis (2015) using equation (4).

## 5 The infinitesimal limit

This general model converges to a normal distribution when we take the infinitesimal limit. We accomplish this by first substituting a Taylor series for the genealogical and mutational distributions in equation (3) (see Appendix A). The infinitesimal limit corresponds to the situation where the effect sizes of mutations become small as the number of loci becomes large. The resulting distribution is multivariate normal where the expected trait value is 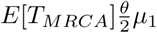, the variance is 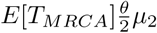, and the covariance between trait values in two individuals *a* and *b* is 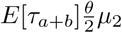. E[*T*_*MRCA*_] is the expected time to the most recent common ancestor in the sample or population. 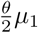 is the rate per unit time per genome that mutational bias shifts the mean. 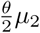 is the rate per unit time per genome that variance accumulates. This limit requires that the products of *L* and moments three and greater of the mutational distribution go to zero as the number of loci becomes large and the effect size per mutation becomes small. This can be thought of as requiring the mutational distribution to not have too heavy of tails. Details of the derivation are given in Appendix A.

Interestingly, the rate of variance accumulation is proportional to the second moment of the mutational distribution instead of the variance. We can see the intuition for this by considering a degenerate distribution where each mutation has the same effect. Here, we would still expect variation among individuals due to differences in the number of mutations each individual receives, even though the variance of the mutational distribution is zero. The variance among individual trait values thus has one component due to differences in the number of mutations and an additional component due to differences in the effects of these mutations. The first component is proportional to the square of the mean mutational effect, while the second is proportional to the mutational variance. Therefore, the sum of the two components is proportional to *m*_2_, the mean squared effect.

Since the trait values are normally distributed, any linear combination of sampled trait values will be as well. This includes the distributions of observable quantities like the differences in trait values from a reference individual or from a sample mean. The distribution of trait differences between individuals is multivariate normal with mean zero and covariance between any pair of trait differences given by

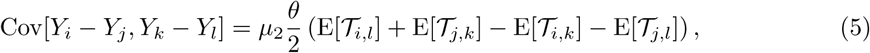

where 𝒯*_i,j_* = 0 if *i* = *j*. Classical theory in quantitative genetics uses a univariate normal distribution of phenotypes in a panmictic population. We can recover this by considering a population of exchangeable individuals. In this case E[𝒯*_i,j_*] = E[𝕋_2,2_] for all pairs *i* ≠ *j*. Individual trait values are then conditionally independent given the mean value in the population and are normally distributed with variance 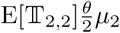.

The normal model in the infinitesimal limit provides additional theoretical justification for studies using normal models to look for differences in selection on quantitative traits between populations (Ovaskainen *et al.*, 2011; Praebel *et al.*, 2013; Robinson *et al.*, 2015). Additionally, equation (5) implies that a covariance matrix based on mean pairwise coalescent times rather than population split times should be used in phylogenetic models of neutral trait evolution.

## 6 Low-mutation-rate approximation

A useful simplification of the model considered so far is to ignore the possibility of more than one mutation per locus. This approximation is reasonable as long as the nucleotide positions affecting the trait are loosely linked throughout the genome. The low-mutation-rate approximation greatly simplifies the mgf of the trait distribution such that it is no longer necessary to know the full form of the mgf of the genealogy:

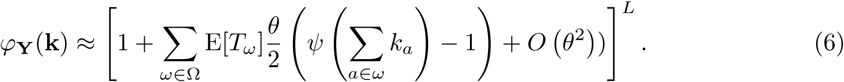

Equation (6) ignores terms that are order two and above in the mutation rate. Conveniently, the mgf of the trait values depends only on the expected length of each branch, whereas equation requires second order and greater moments of branch lengths (e.g. E[*T*_*ω*1_ *T*_*ω*2_]). We can use equation (6) to express moments of the trait distribution in terms of expected branch lengths calculated from coalescent models.

## 7 Moments of the trait distribution

For most population genetic models and for sample sizes greater than three, the recursive nature of the trait distribution mgf makes it computationally unfeasible to solve under general parameter values. However, it is not necessary to have an expression for the full mgf in order to derive moments of the trait distribution in terms of moments of branch lengths and mutational effects. Under the low mutation rate approximation moments can be calculated by differentiating equation (6). Even without making this approximation, moments can be calculated by taking Taylor expansions in equation (2) and only considering terms contributing to the desired moment’s order. We implemented a symbolic math program to calculate trait moments using this procedure, and the details are given in Appendix B. As the normal distribution is completely defined by its first two moments, the extent to which a trait distribution deviates from normality can be measured by the extent to which its moments deviate from those of a normal distribution with the same mean and variance.

We have so far considered the distribution of a trait value *Y*_*a*_ over evolutionary realizations. The expectation of *Y*_*a*_ is 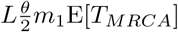, and the variance is 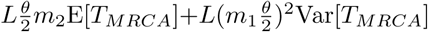. Although simple to derive using computer algebra, expressions for the higher central moments of *Y*_*a*_ are complicated even under the low mutation rate approximation, and there is not much to be gained by showing them here.

However, in a given evolutionary realization there will be a distribution of trait values in the population. The population-level trait distribution can also be described by its moments, but since this distribution is random the moments at the population level are also random quantities. Since *Y*_*a*_ is relative to a value that is not directly observed, the expected population-level moments offer more insight. In particular, we are interested in how the trait distribution at the population level might deviate from normality. Schraiber and Landis (2015) computed the expected first four central moments of a constant-size population. We derive the same expectations under an arbitrary demographic history,

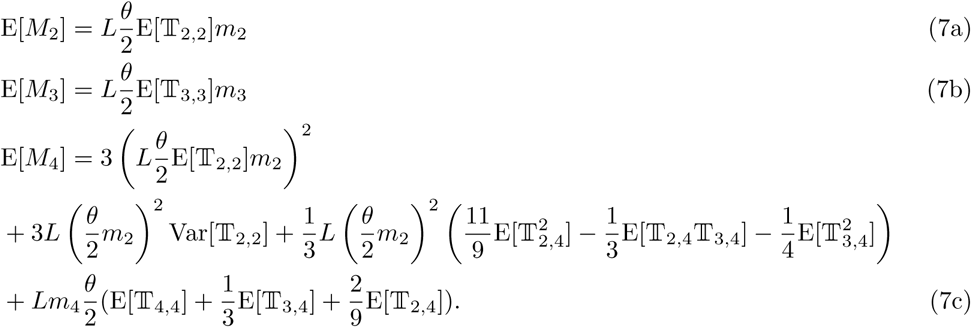

The normal limit corresponds to equation (7b) and the second two lines of equation (7c) going to zero. To give insight into the expressions in equation (7), moment calculations done by hand under the low mutation rate approximation are presented in Appendix D.

Equation (7a) gives the expected trait variance in the population. However, the variance will vary over realizations of the evolutionary process. The variation in the population variance depends on the sparsity of the trait and the number of causal loci. The variation in the variance can be quantified using its coefficient of variation (CVV), the standard deviation of the variance divided by its expectation. For a constant-size, panmictic population

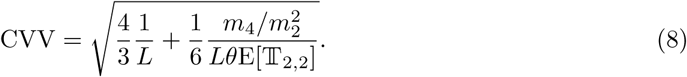

Equation (8) shows a contribution due to linked mutations occurring at a single locus 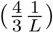 and a contribution due to sparsity and mutation 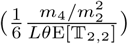. Even when the sparsity is low, i.e., when a large number of variants affect the trait, if the trait is only controlled by a single locus there will be considerable variation in the population variance 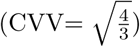. On the other hand, the CVV of a sparse trait controlled by many loci will depend on the ratio of the fourth and squared second non-central moments of the mutational distribution 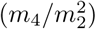.

## 8 Comparison to normal distribution

Deviations of the population distribution from normality depend on the distribution of coalescent times and the genetic parameters. One natural way to quantify a distribution’s deviation from normality is through its kurtosis. The kurtosis, defined as 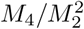, measures the tendency of a distribution to produce outliers (Westfall, 2014). However, since the kurtosis is a ratio of two random variables, its expectation is challenging to calculate. Rather than attempt this calculation, we compare the expected fourth central moment itself to that expected under normality. This approach qualitatively identifies the factors influencing deviations from normality.

Using the low mutation rate approximation, the ratio of the expected fourth moment to that under normality simplifies to

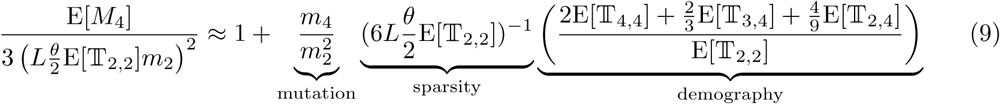

The expected excess in *M*_4_, relative to normality, increases with sparsity and with 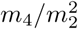. 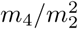 is equal to the kurtosis of the mutational effect distribution when mutations are unbiased and reflects its propensity to produce large effect mutations. The excess depends on demography through the factor 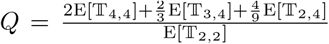. The extent to which demography increases or decreases deviations from normality can be investigated by calculating *Q* in different models. In a constant-size and panmictic population *Q* is equal to one. In a population where lineages are exchangeable, E[𝕋*_k,n_*] can be calculated numerically using expressions from Griffiths and Tavaré (1998) or Polanski and Kimmel (2003). Values of *Q* in an exponentially growing population are shown in Figure 2A. Holding sparsity and the mutational distribution constant, a population which underwent exponential growth will have a greater expected deviation from normality in its trait distribution. Another example demography is a population that goes through a step change at some point in the past. Figure 2B shows that when the population size increased at some point in the past, *Q* is increased similarly to the exponential growth scenario. When the population experiences a bottleneck *Q* is decreased below one. Additionally, *Q* appears more sensitive to population growth than to bottlenecks.

**Figure 2:**
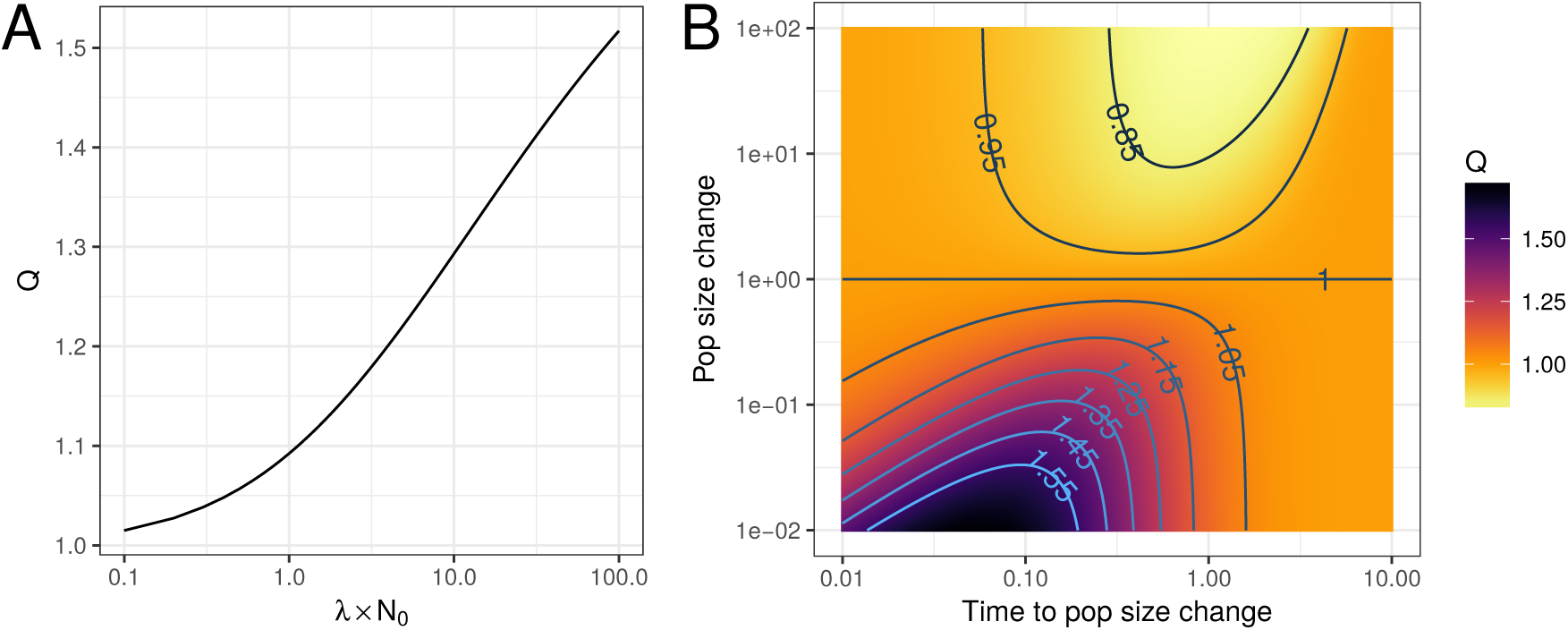
The effects of demography on deviations of the expected fourth central moment of the population trait distribution from normality. ***Q*** measures the effect due to demography on the expected fourth central moment (equation (9)). (**A**): *Q* increases as the exponential growth rate increases relative to the current population size. *λ* is the growth rate and *N*_0_ is the initial effective population size. (**B**): *Q* values when the population undergoes an instantaneous step change at some point in the past. The time and magnitude of this change are given in units of the current effective population size. *Q* increases when the population grows and decreases when it declines.

As a concrete example, we can consider the differences in the expected fourth moment produced by different demographic histories in different human populations. In the demographic model fit by Tennessen *et al.* (2012), the generic European population experiences a bottleneck associated with out-of-Africa and recent growth while the generic African population experiences a more stable history also with recent growth. Differences in demographic history between the two populations has resulted in a lower heterozygosity in European populations due to the out-of-Africa bottleneck (Yu *et al.*, 2002).

For a given sparsity, the African population model predicts a smaller deviation from normality than the European model (Figure 3). The expected fourth moment in constant-size populations with the same heterozygosity as the African and European models is lower for the African model and higher for the European model. This is because the African model is dominated by population growth that leads to a *Q* greater than one, while the European model is dominated by a bottleneck event that leads to a *Q* less than one (Figure 2). However, differences due to demography are small and deviations from normality are mostly driven by differences in heterozygosity at causal loci.

**Figure 3:**
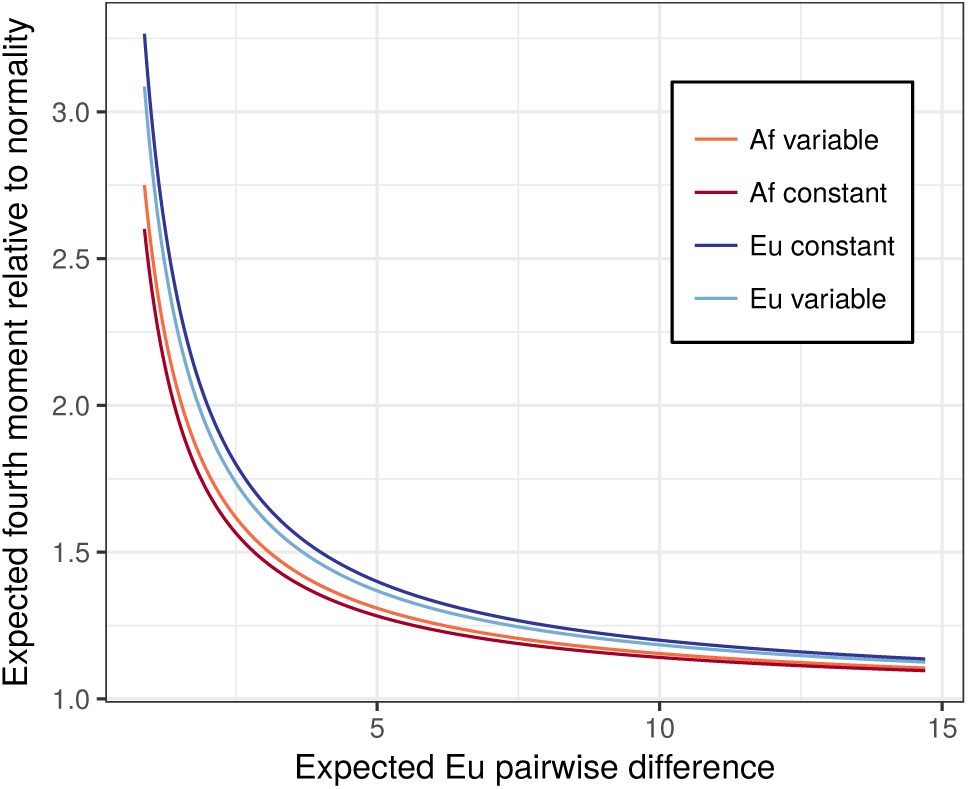
A comparison between the expected fourth moment at different levels of sparsity in the African and European demographic models fit by Tennessen *et al.* (2012). Trait sparsity is varied by changing the expected number pairwise differences at sites affecting the trait in the European model. The mutational kurtosis is set to six. The darker lines show the predicted relationship for populations with the same heterozygosity as the European and African models but with constant size.

Even though recombination is not included in the model, we can form an idea about how linkage might impact deviations from normality. Line two of equation (7c) corresponds to the contribution from two mutations occurring at a single locus. The first quantity indicates that the expectation of the fourth moment increases with the variance of the pairwise coalescence time. The second part does not have a clear interpretation. If the sum of these two terms is positive this agrees with the intuition that linkage disequilibrium increases deviations from normality by reducing the effective number of independent loci.

Simulations of the kurtosis itself show substantial variance, with about a quarter of simulated populations having a value less than three (that of a normal distribution) even as the mean kurtosis increases to almost nine (Appendix C). This high variance in the kurtosis is likely due to a high variance in both the trait variance and fourth moment. This, along with the fact that deviations from the infinitesimal model inflate the fourth moment (equation (9)), leads to a situation where the kurtosis increases with trait sparsity but the variance is high across evolutionary realizations.

## 9 Trait divergence in structured populations

To demonstrate the utility of the normal distribution arising in the infinitesimal limit (equation 5), we derive a simple procedure for simulating from null distribution for the divergence between groups in structured populations. A common way to quantify the divergence in trait values between groups is *Q*_*ST*_, defined as the variance between the group means divided by the total variance in the population (Spitze, 1993). In a haploid model, 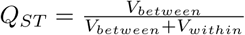, where

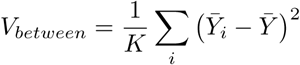

and

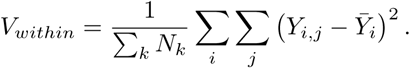

Here, *Y*_*i,j*_ is the trait value of individual *j* in population *i, K* is the number of subpopulations, and *N*_*k*_ is the size of subpopulation *k*.

In the normal model, all *Y*_*i,j*_ are normally distributed. Therefore, 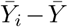and 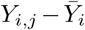 are also normally distributed. When population sizes are large, individual deviations from population means are nearly uncorrelated, as are *V*_*between*_ and *V*_*within*_. *V*_*within*_ is nearly constant across evolutionary realizations and is equal to ∑*_k_ N*_*k*_*E*[𝒯*_k,k_*]*/* ∑*_k_ N_k_* because the within population variances are approximately uncorrelated and their variances are order 1*/N_k_*. While we do not have an explicit form for the density function of the between-group variance, we can simulate from its distribution by drawing a vector of 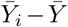 values from a multivariate normal distribution with mean zero and with a covariance matrix whose element between populations *a* and *b* is

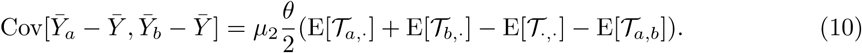

To simulate *Q*_*ST*_ values we do not need to know *µ*_2_ or 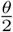 because the scale of the trait variance cancels in the *Q*_*ST*_ ratio. Therefore, all that is necessary to simulate from the null distribution of *Q*_*ST*_ is to have estimates of the expected coalescent time within and between populations. No further information about the history or structure of the population is needed. Since only the relative coalescent times matter it is also not necessary to scale these estimates to units of years or generations using a mutation rate.

This procedure for simulating from the null distribution of *Q*_*ST*_ could be useful in testing whether an observed *Q*_*ST*_ is unlikely under neutrality. Current goodness-of-fit tests either compare *Q*_*ST*_ to an empirical distribution of *F*_*ST*_ values or to a *χ*^2^ distribution (Leinonen *et al.*, 2013). In the second case, an identical distribution to that developed by Lewontin and Krakauer (1973) is used as the null distribution for *Q*_*ST*_. The *χ*^2^ testing procedure was suggested by Whitlock and Guillaume (2009) and is implemented in the program *QstFstComp* (Gilbert and Whitlock, 2015). The Lewontin-Krakauer (LK) distribution assumes independence between subpopulations and provides a good approximation in populations without spatial structure, such as in a symmetric island model of migration (Figure 4). When subpopulations are strongly correlated, such as in a one-dimensional stepping-stone model of population structure, the LK distribution is a poor approximation. Even when the distributions appear qualitatively similar, there are substantial differences in tail probabilities (Figure 5).

**Figure 4:**
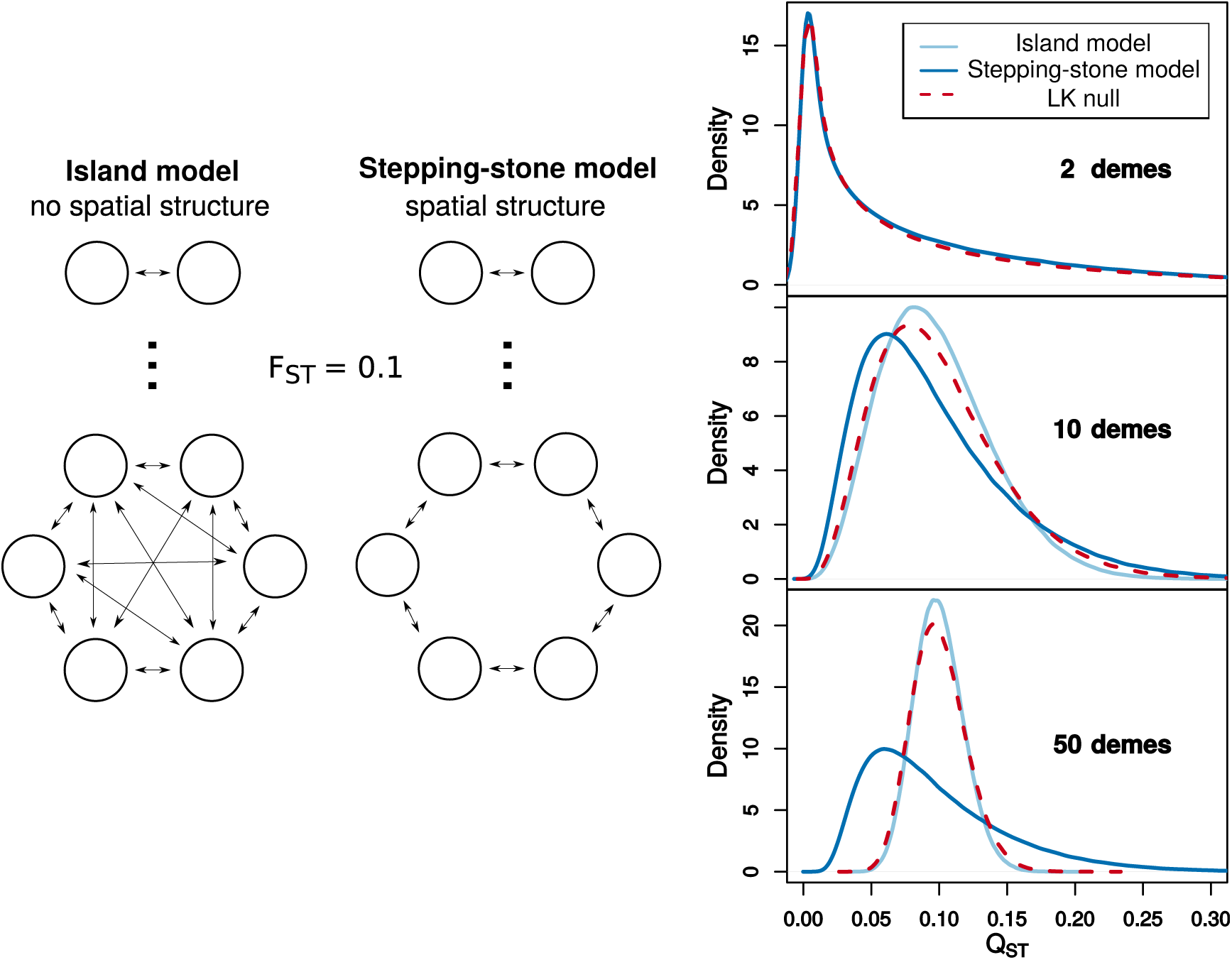
A comparison of neutral sampling distributions for *Q*_*ST*_ at the population level. The Lewontin-Krakauer (LK) distribution for *Q*_*ST*_ is compared to the null distribution in the infinitesimal limit in a case with (the stepping-stone model) and a case without (the island model) spatial structure. The case with no spatial structure assumes the migration rate is equal between all subpopulations, and the case with spatial structure arranges subpopulations in a ring with migration only between neighboring subpopulations. Migration rates and subpopulation sizes are all equal and are set such that *F*_*ST*_ = 0.1 even as the number of subpopulations is increased (Slatkin, 1991). *Q*_*ST*_ values for these models are simulated by drawing vectors from a multivariate normal distribution parameterized using the expected pairwise coalescence times given by (Slatkin, 1991). Under the LK distribution *Q*_*ST*_ is distributed as *F*_*ST*_ /(*n*_*d*_ − 1) times a chi-square distribution with *n*_*d*_ − 1 degrees of freedom 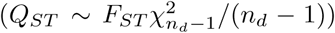, where *n*_*d*_ is the number of subpopulations. Vertical lines show the mean *Q*_*ST*_ under the different null distributions. The mean *Q*_*ST*_ values for the two normal models are nearly identical. Discordance between these lines and that for the LK distribution illustrates that *Q*_*ST*_ ≠ *F*_*ST*_. The neutral distributions for *Q*_*ST*_ become increasingly different as the number of subpopulations over which the genetic divergence is spread increases.

**Figure 5:**
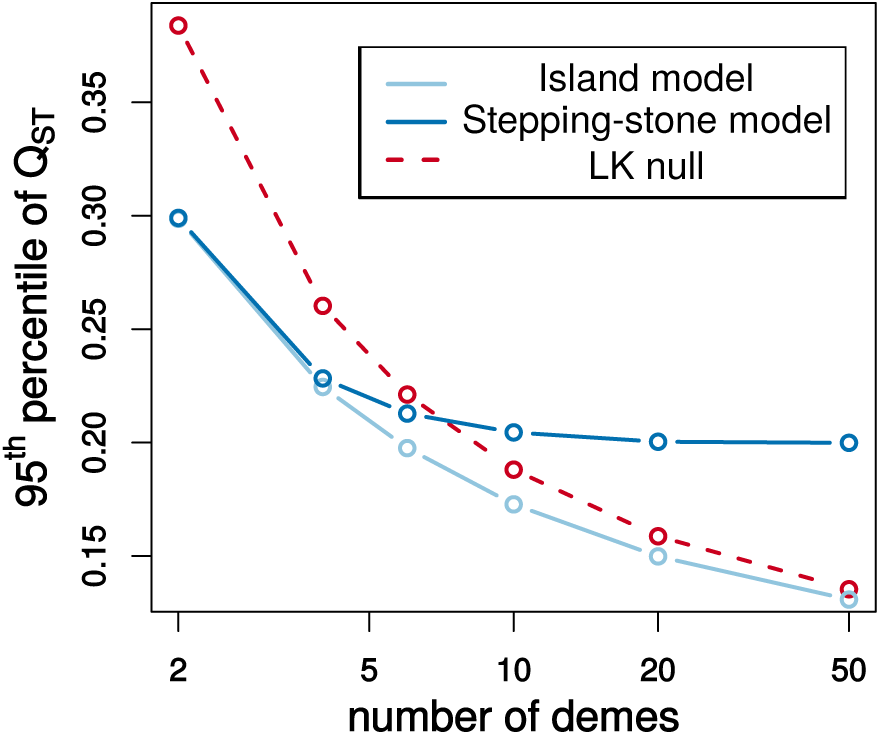
Differences in the 95^th^ percentile of different neutral sampling distributions for *Q*_*ST*_ at the population level. The Lewontin-Krakauer (LK) distribution is compared to neutral null distributions for structured populations with and without a spatial component using the multivariate normal model that arises in the infinitesimal limit.

Nearly identical issues to these were raised with the LK test at the time of its publishing (Nei and Maruyama, 1975; Robertson, 1975). However, while the neutral distribution for *F*_*ST*_ depends on the precise details of population structure and history, the distribution of *Q*_*ST*_ only depends on the set of within and between subpopulation coalescence times. The point here is not that the LK distribution is particularly bad, but rather that an improved neutral distribution can be obtained using the normal model arising in the infinitesimal limit. The neutral distribution described here is similar to the extension of the LK test developed by Bonhomme *et al.* (2010) to account for the correlation structure between subpopulations. The Bonhomme *et al.* (2010) method treats allele frequencies as multivariate normal with covariance matrix parameterized by coancestry coefficients. Ovaskainen *et al.* (2011) use a normal model similar to that found in the infinitesimal limit here, but the covariance matrix is also based on coancestry coefficients. When phenotypic and genetic divergence is mostly driven by changes in allele frequency, the coalescent and coancestry based models should be very similar. However, the coalescent model is ultimately preferable since it is the correct null model at any scale of population divergence in the infinitesimal limit. When only allele frequency data are available, a coancestry model is the only option, but it is still better to model variable levels of shared ancestry between populations than to use a single value of *F*_*ST*_.

Additionally, treating Q_ST_ as a random variable lets us reexamine the classic result in evolutionary quantitative genetics that *Q*_*ST*_ = *F*_*ST*_ (Whitlock, 1999). *F*_*ST*_, in this context, refers to a parameter of the population. In particular, 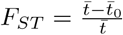, where 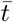 is the expected coalescent time for two loci sampled at random from the entire population and 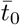 is the expected coalescent time for two loci sampled within a subpopulation (Slatkin, 1991). This value is constant over realizations of the evolutionary process. *Q*_*ST*_ can only refer to either state of the population or to an estimate of this state. As shown above, *Q*_*ST*_, as a state of the population, varies across evolutionary realizations. Thus, there is no sense in which *Q*_*ST*_ can be defined as a constant parameter in the way that *F*_*ST*_ can. The expectation of *V*_*between*_ is 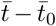, and the expectation of *V*_*within*_ is 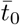. 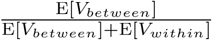 is equal to *F*_*ST*_, but due to Jensen’s inequality, the expectation of this ratio (E[*Q*_*ST*_]) is always less than *F*_*ST*_.

## 10 Inferring genetic parameters

Schraiber and Landis (2015) suggested it might be possible to infer the shape of the distribution of mutational effects through its moments for sparse traits whose distributions deviate from normality. Using the expressions for the expected moments of the trait value distribution in equation (7), we could try and design a method of moments estimator for the moments of the mutational distribution. If 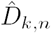 is an estimator of 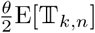, the system of equations for the first three central moments under the low mutation rate approximation is

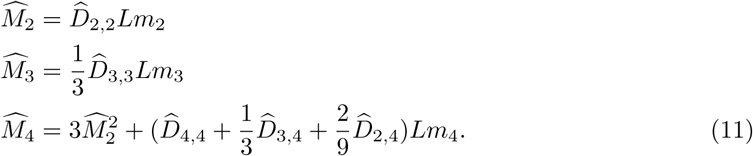

From equation (11) we can see that the moments of the trait distribution only enter through products with the number of loci potentially affecting the trait (*Lm*_*i*_). If these products were estimated as 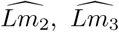, and 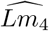, it would be possible to estimate the ratios *m*_2_*/m*_4_ and *m*_3_*/m*_4_ of the moments of the mutational distribution. These ratios are meaningless on their own because any value could be obtained by changing the scale on which the trait is measured. The quantity that is identifiable in this system of equation is the compound parameter 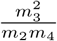. This quantity reflects something about the mutational bias relative to the spread of the distribution. However, it is likely not possible to distinguish sparsity from skewed or heavy-tailed distributions of mutational effects.

## 11 Discussion

Neutral models of quantitative trait evolution are important for establishing a baseline against which to test for selection. Schraiber and Landis (2015) recently analyzed a neutral model of trait evolution that made few assumptions about the number of loci potentially affecting the trait and the distribution of mutational effects at these loci. However, they only derived results for constant-size, panmictic populations. We extend their results to populations with arbitrary distribution of coalescent times and therefore varying demographies and population structures. As their key result, Schraiber and Landis (2015) derived the characteristic function of the distribution of trait values in a sample. In this paper, we instead work with the moment generating function, but the two approaches are interchangeable as long as the mgf for the distribution of mutational effects exists (which it will if the moments of the mutational distribution are all finite). Our main result, given in equation (4), shows that the characteristic function from Schraiber and Landis (2015) is a special outcome of a general procedure whereby the moment generation function for a trait distribution can be obtained by making a simple substitution into the moment generating function for a distribution over genealogies. The moment generating functions for many demographies of interest and for sample sizes above three are sufficiently complex that solving for them is impractical (Lohse *et al.*, 2011). However, progress can still be made by using Taylor expansions to write moments of the trait distribution in terms of moments of the genealogical and mutational distributions.

This result extends previous work using coalescent theory to investigate neutral models of quantitative traits (Whitlock, 1999; Schraiber and Landis, 2015). Ours is the most general model yet analyzed. As a natural first step, we show that the infinitesimal limit suggested by

Fisher (1918) leads to a model where phenotypes are normally distributed as the number of loci becomes large and the variance of effect sizes becomes small. In the limiting distribution, the variance of the difference in trait values between two individual is proportional to the expected pairwise coalescent time between them, and the covariance between a pair of differences more generally is Cov[*Y*_*a*_ − *Y*_*b*_, *Y*_*c*_ − *Y*_*d*_] *∝* E[*𝒯_a,d_*] + E[*𝒯_b,c_*] − E[*𝒯_a,c_*] − E[*𝒯_b,d_*]. The resulting covariance matrix completely specifies the neutral distribution under the infinitesimal model. This is similar to classic models in evolutionary quantitative genetics considering the neutral divergence of trait values after population splits (Lande, 1976; Lynch, 1989) but holds regardless of the precise details of population structure and history. Schraiber and Landis (2015) derive essentially the same distribution using a central limit theorem argument.

Mendes *et al.* (2018) point out several problems caused by incomplete lineage sorting when species split times are used to generate the covariance matrix in phylogenetic comparative methods. Since the normal model arising in the infinitesimal limit is explicitly based on the coalescent, it is not subject to these problems. A matrix based on pairwise coalescence times rather than a species tree based on population split times takes into account the effects of all lineages at causal loci, even those that do not follow the species topology. The covariances specified by equation (5) could also be used to generate within-species contrasts in a similar manner to the method suggested by Felsenstein (2002).

We compared sparse traits to the normal model by calculating how the first four expected central moments different from those expected under normality. This showed how demography and genetics separately influence the expected deviation from normality. For a fixed expected trait sparsity, population growth produces greater deviations in the fourth central moment while population bottlenecks produce lower deviations (Figures 2). However, for realistic demographic scenarios, we find that the effects attributable to demography are small (Figure 3). We only analyzed cases where the individuals in the population were exchangeable, but adding population structure would increase deviations from normality as drifting trait means between subpopulations will yield a multimodal distribution.

We next apply the above theory to two simple problems where a coalescent perspective on the neutral distribution of a quantitative trait provides useful intuition. The first of these is the question of the appropriate null distribution for *Q*_*ST*_ at the population level. We show how the null distribution under the normal model can be easily simulated from by taking advantage of the theory presented here, providing a much better approximation when populations are correlated than previous approaches (Whitlock and Guillaume, 2009) (Figure 4). In the second we show that the shape of the mutational distribution is largely confounded with the number of loci affecting the trait, with only one compound parameter identifiable. This makes it unlikely that mutational parameters could be inferred from trait values sampled from a population.

Even though we have broadened the model space for neutral traits, many features of real populations have not yet been incorporated. Linkage between loci is a particular concern as there is substantial linkage disequilibrium between causal loci (Bulik-Sullivan *et al.*, 2015). Lohse *et al.* (2011) derived the form of the moment generating function for linked loci and future work will attempt to incorporate this using equation (4). In particular, it will be important for future work to show how this affects the distribution in the infinitesimal limit. Diploidy, dominance, and epistasis have also been ignored thus far. The qualitative effects described here should hold under diploidy, but having trait values within individuals summed over loci from two copies of the genome will decrease deviations from normality. Dominance will also tend to produce a normal distribution as the effects are independent between loci, but future work is needed to examine how this interacts with the distribution of genealogies to affect the trait distribution.

Barton *et al.* (2017) recently performed a deep mathematical investigation of a more formal “infinitesimal model” that is subtly different from the infinitesimal limit considered here. They proved conditions under which the trait values of offspring within a family are normally distributed with variance independent of the parental trait values conditional on the pedigree and segregation variance in the base population. Interestingly, they found the normality for offspring trait values still holds under some forms of pairwise epistasis that are not too extreme. This implies it may be possible to include epistasis in the infinitesimal limit considered here. It would be important to know how epistasis affects the neutral divergence of trait between populations and species.

Although GWAS of many traits have shown them to be controlled by large numbers of loci (Boyle *et al.*, 2017), this will not necessarily be the case for every trait of interest to biologists. It has been suggested, for instance, that gene expression levels have a sparse genetic architecture (Wheeler *et al.*, 2016). Since there is much interest in testing whether natural selection has acted on gene expression levels (Whitehead and Crawford, 2006; Gilad *et al.*, 2006; Yang *et al.*, 2017), well-calibrated goodness-of-fit tests will need to take into account the complications that arise when trait distributions deviate from normality (Khaitovich *et al.*, 2005). Direct measurements of mutational distributions (Gruber *et al.*, 2012; Metzger *et al.*, 2016) could aid in such a calibration. Finally, equation (7) suggests a means to determine whether sparsity impacts trait distributions by examining populations with different demographic histories. Populations with a greater 𝕋_3,3_ to 𝕋_2,2_ ratio are also expected to show a greater *M*_3_ to *M*_2_ ratio and an equivalent relationship would be expected for the fourth moment. Because gene expression studies generally measure a large number of traits, observing such a trend on average could be a sign of deviations from normality. Generally, more work is needed to develop neutrality tests that are robust to the details of genetics and population history, and to investigate whether any information about mutation can be uncovered by statistical models that share information across multiple measured traits.

## Acknowledgements

We thank John Novembre, Daniel Rice, and members of the Novembre lab for helpful discussions. We would also like to thank Josh Schraiber, Daniel Rice, John Novembre, Maryn Carlson, Hussein Al-Asadi, and Joe Marcus for their comments on the manuscript. This work was supported by an NSF GRFP to the author and NIH grant GM108805 to John Novembre.

## A A central limit theorem in the infinitesimal limit

Recall that the moment generating function for the distribution of trait values from a single locus is

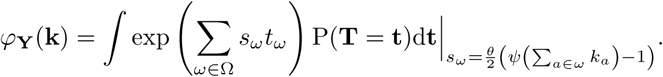

If we substitute in the Taylor series expansions for the moment generating function of the trait value distribution we get

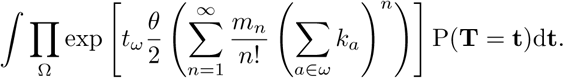

If we then write the Taylor series of each exponential function we get

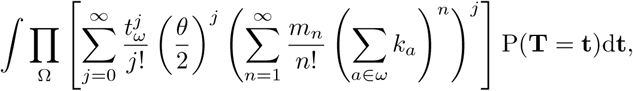

which is equivalent to

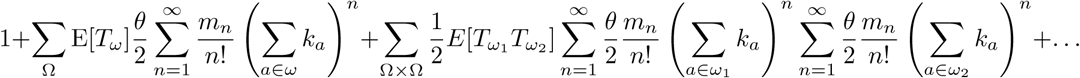

This is raised to the power *L* for a trait controlled by *L* loci. We want the limit as the number of loci increases while the size of mutational decreases. This can be expressed by the limits *Lm*_1_ → *µ*_1_, *Lm*_2_ → *µ*_2_ and *Lm*_*i*_ → 0 for *i >* 2 as *L* → *∞*. Knowing we will not be retaining *m*3 and above we can rewrite the mgf as

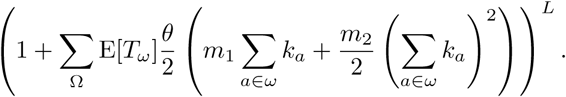

The result of taking these limits is

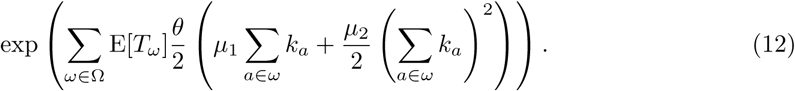

This is multivariate normal distribution with mean equal to 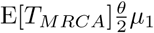, variance equal to 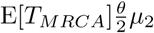, and covariance between *Y*_*a*_ and *Y*_*b*_ equal to 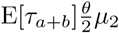. This can be seen from equation (12) by noting that in the mgf for a multivariate normal distribution the coefficient in the exponential of *k*_*a*_ is the mean of *Y*_*a*_ and the coefficient of *k*_*a*_*k*_*b*_ is 2Cov[*Y*_*a*_, *Y*_*b*_] if *a* ≠ *b* and Var[*Y*_*a*_] if *a* = *b*.

These *Y* values are not directly observed because in the theory presented here they are measured as the sum of differences since the *T*_*MRCA*_s at the causal loci affecting the trait. Rather, differences between individual trait values are what analyses would be based on. Since the *Y* are normally distributed differences between trait values such as *Y*_*a*_ − *Y*_*b*_ will be as well.

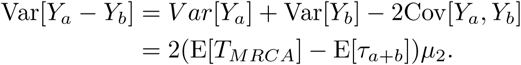

This also tells us that the distribution of trait differences will be the same in the infinitesimal limit with and without a low mutation rate approximation.

A much simpler heuristic derivation of the limiting normal distribution can be done by calculating the variance and covariance at a single locus. This derivation is very similar to that done by Schraiber and Landis (2015). Using the law of total variance we can write

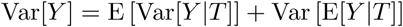

The variance conditional on *T* can be calculated again using the law of total variance and conditioning on the number of mutation at the locus.

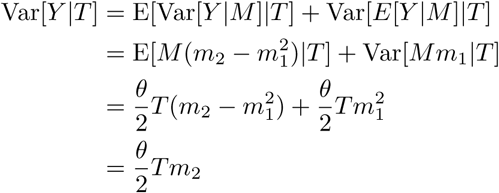

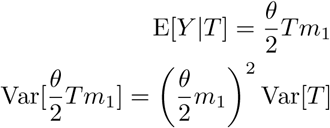

Therefore we have

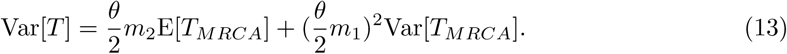

The same procedure can be done for the covariance.

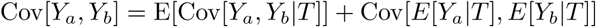

We can break *Y*_*a*_ and *Y*_*b*_ into a shared part, *Y*_*S*_ and unshared parts for each, *Y*_*δa*_ and *Y*_*δb*_.

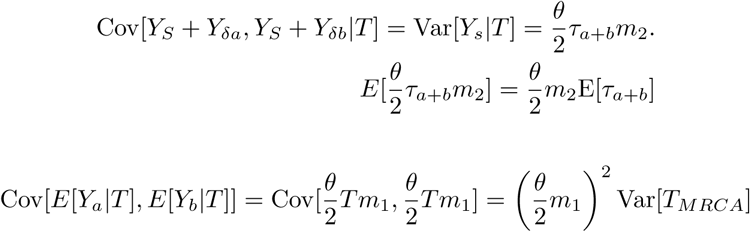

Therefore we have

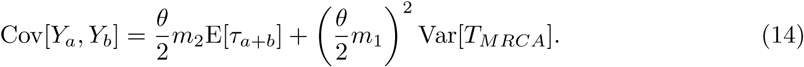

The terms proportional to the variance of the *T*_*MRCA*_ disappear because 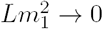 as 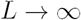.

## B Automatic moment derivations

Moments of the trait distribution can be calculated by differentiating equation (3). To automate the process, symbolic math programs were written using sympy (Meurer *et al.*, 2017). To derive moments for an arbitrary distribution of coalescent times we take Taylor series of the mgf of the mutational distribution to get

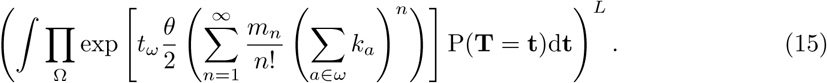

We then also Taylor expand the exponential function appearing in this integral to get

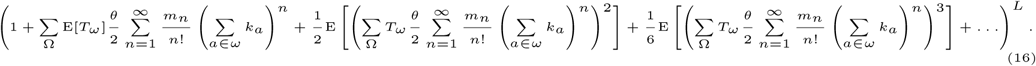

Of course, the infinite sums appearing in this expression pose a problem. To calculate a particular moment, we only consider terms that will contribute terms of that moment’s order when equation (16) is differentiated and zero is substituted for all dummy variables. For example, to calculate *E*[*Y*_*a*_*Y*_*b*_*Y*_*c*_] we only want terms that contain an order three product of *k* when the polynomial in equation (16) is expanded. For a moment of order *m*, we only need consider terms of size *m* and less in the series expansion of the exponential, and within the expansions of the mutational mgf we only need consider terms up to order *m − m_t_* + 1 where *m*_*t*_ is the order of the exponential term being considered.

## C Kurtosis simulations

**Figure S1:**
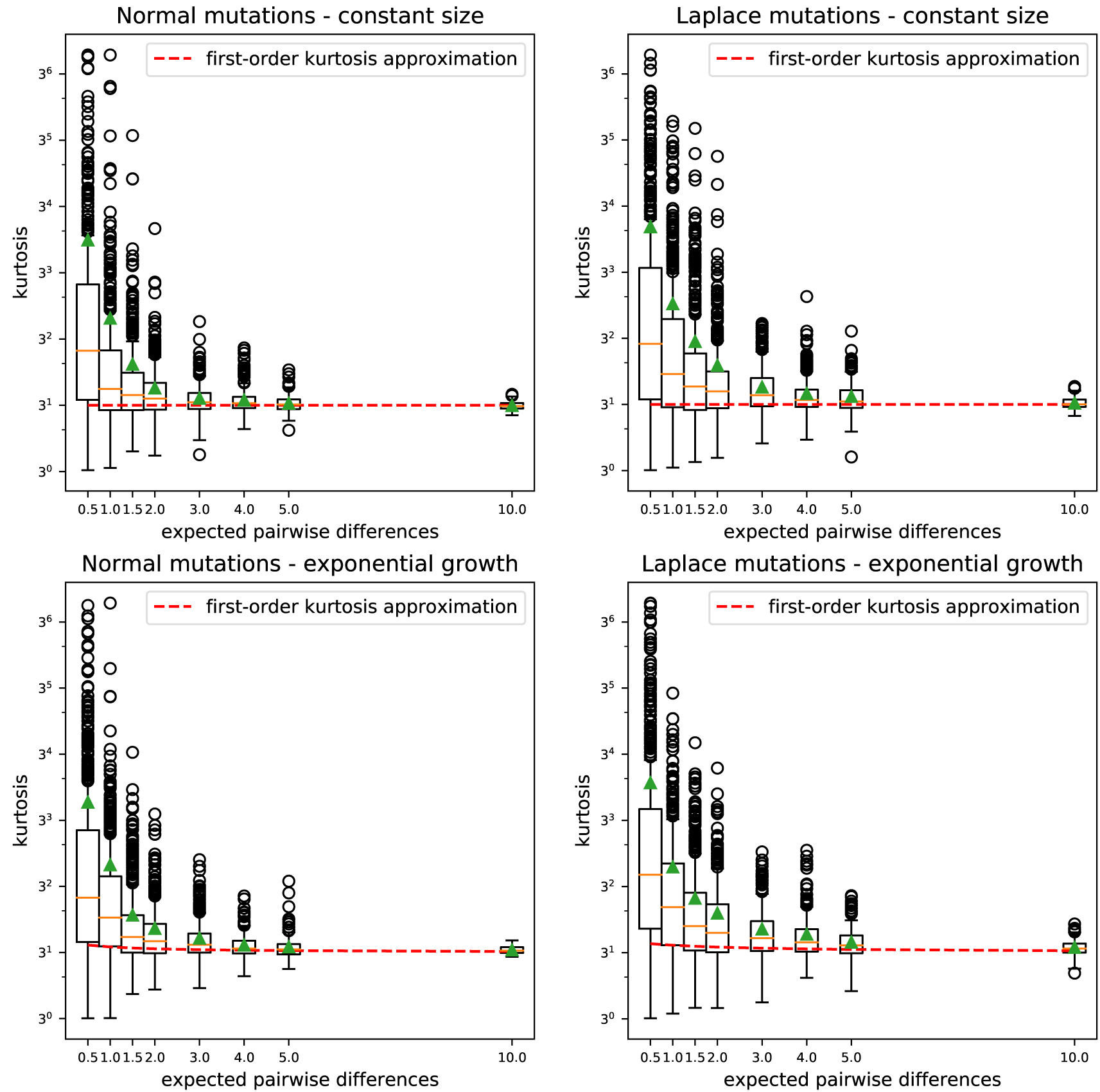
The distribution of the population kurtosis under different levels of sparsity, mutational kernels, and demographies. The sparsity is varied by changing the expected number of pairwise differences at sites affecting the trait. Normal and Laplace distributions of mutational effects are compared. A constant size population is compared to an exponential growth scenario with growth rate equal to the reciprocal of the final effective population size. Green triangles denote the mean kurtosis. The dashed red lines give the first order approximation to the expected kurtosis given in equation 17.

Entire populations were simulated using msprime (Kelleher *et al.*, 2015) and mutations were assigned effects from a standard normal or Laplace distribution. The effective population size and mutation rate were kept constant and the expected number of pairwise difference was increased by increasing the number of loci affecting the trait.

A first order approximation to the kurtosis is

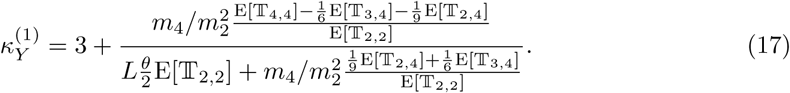

Although this expression suggests that the expected *κ*_*Y*_ will be greater than under normality when external branches are longer 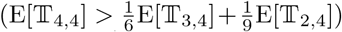 and less than under normality when they are longer 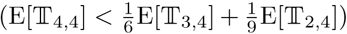, simulations show that the approximation is actually quite poor (Figure S1).

## D Central moment derivations

We can use the low mutation rate approximation to the moment generating function to calculate moments of the distribution of trait vales. We will start by calculating the first and second moments. We start, as we did in deriving the normal distribution, by substituting the Taylor series of the mutational mgf.

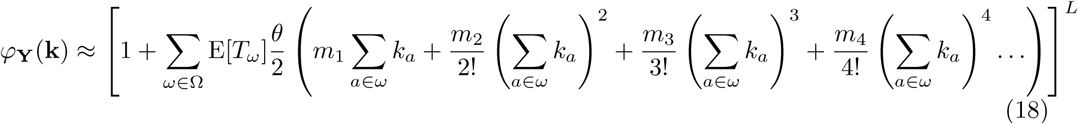

We can expand this out using multinomial coefficients to get

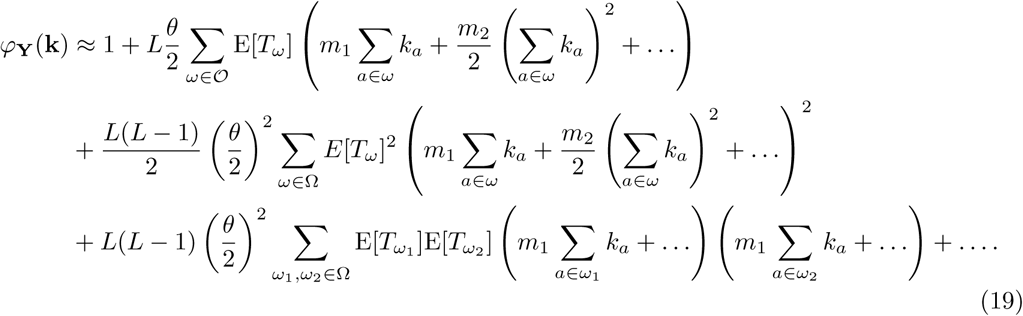

The first coefficient is 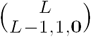, the second is 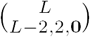, and the third is 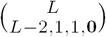. To calculate the moments of this distribution one takes the partial derivatives of the mgf and sets the dummy variables to zero.

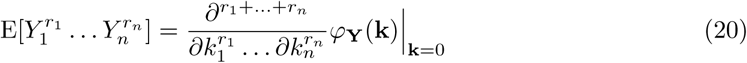

Using this to calculate the first moment of the trait distribution we get

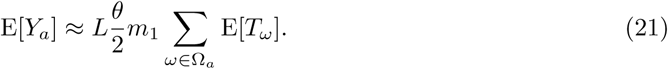

The second moment is more complicated because there are 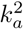 terms in all three lines of equation

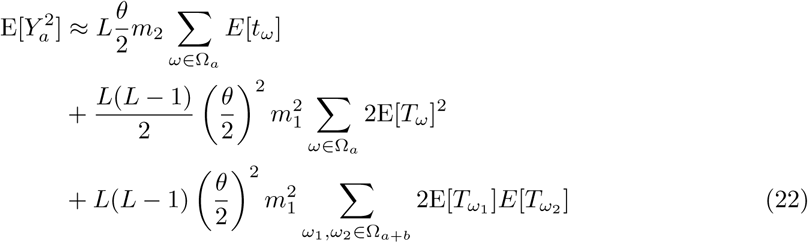

Terms with 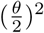 are kept because they also include a second order term of *L* in front of them. We can now calculated the variance using Var[*Y*] = E[*Y*^2^] − E[*Y*]^2^. The squared first moment can be written as

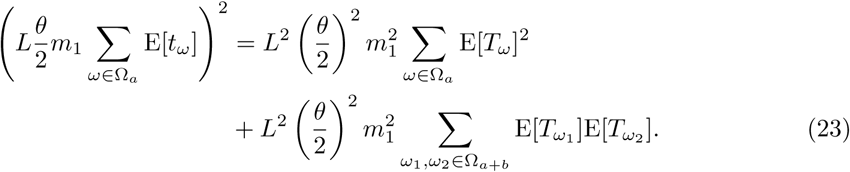

Subtracting this from the second moment gives

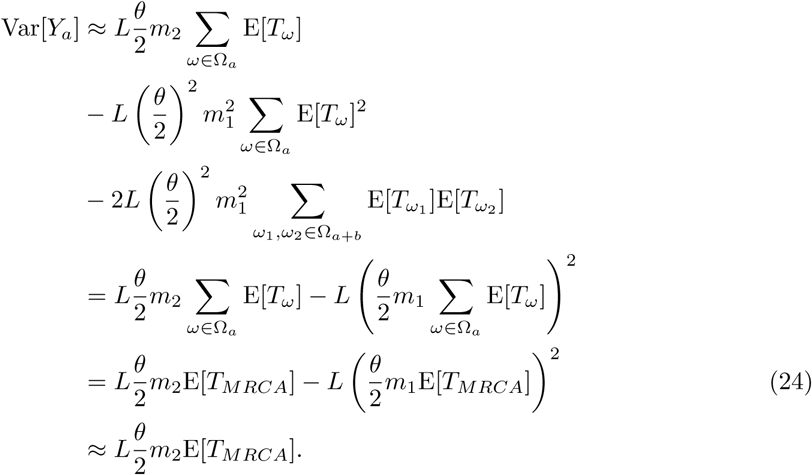

Due to the large number of terms we only derive the fourth moment of the trait value distribution for the case when the mean mutational effect is zero. The terms of (18) appearing in the fourth moment after we apply (20) are

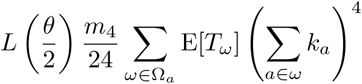

for the fourth moment along one branch,

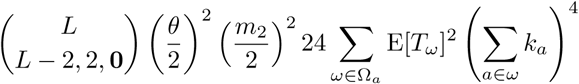

for the second moment of the same branch chosen twice, and

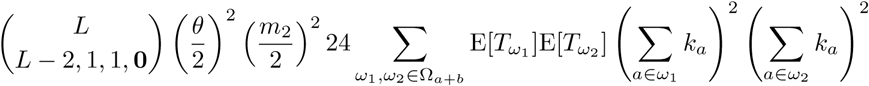

for the second moments on two different branches. Taking the fourth derivatives of these in terms of the desired branch we get

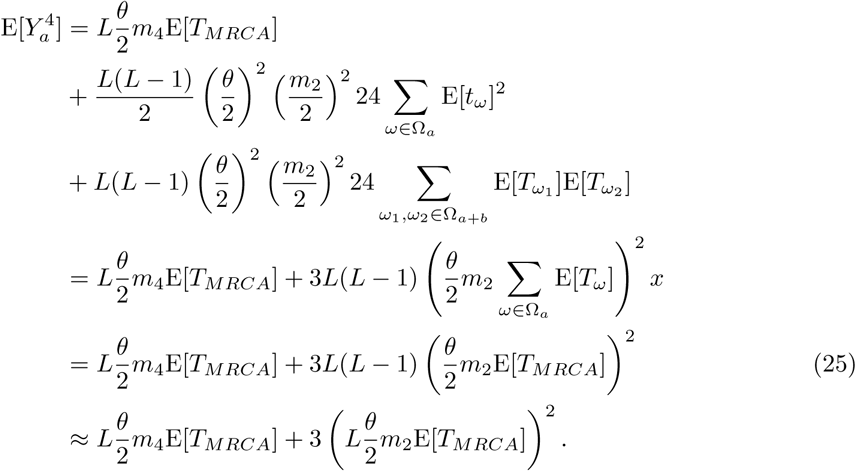

The variance and the fourth moment derived in equation (24) and (25) can be used to derive the kurtosis of *Y*_*a*_. The kurtosis of a random variable *X* is defined as

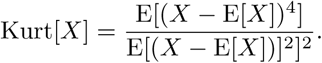

This is the fourth central moment divided by the variance. It is possible to calculate the kurtosis of a single trait value over evolutionary realization. For ease of calculation, we’ll examine this in the case where the mean mutation effect (and therefore trait value) is zero. If we plug (24) and (25) into the expression for the kurtosis we get

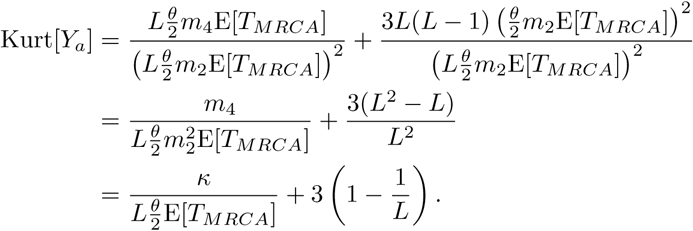

We also calculate some additional moments that have less clear interpretations but which are useful when calculating the expected fourth central moment in the population. The first of these is 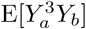. The terms of (18) appearing in this are

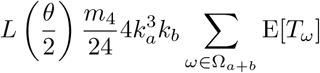

and

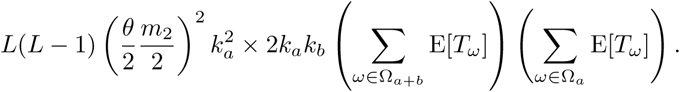

This ultimately gives

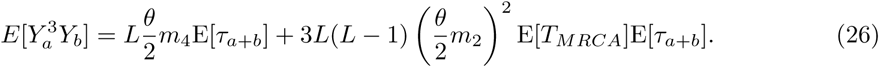

The next fourth moment of interest is 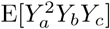. The terms of (18) are

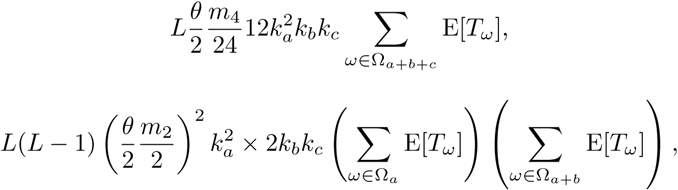

and

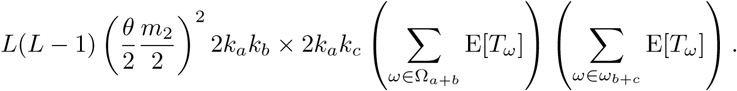

Taking the appropriate derivatives of these gives

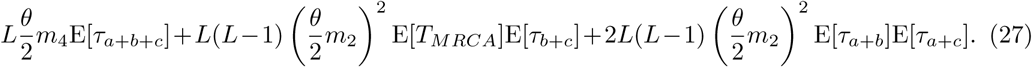

Individuals in the population are exchangeable as long as it is not structured. The pairwise expected shared branch lengths are in that case all equal and we can write (27) as

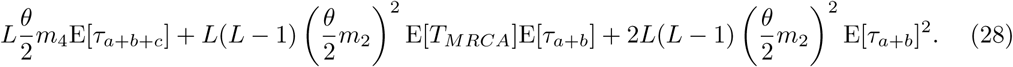

The final moment we’ll look at is E[*Y*_*a*_*Y*_*b*_*Y*_*c*_*Y*_*d*_] which has relevant terms

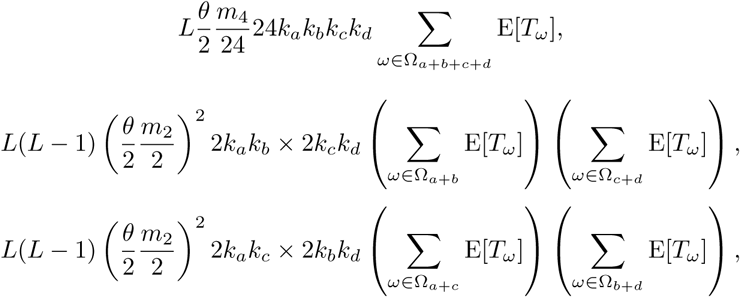

and

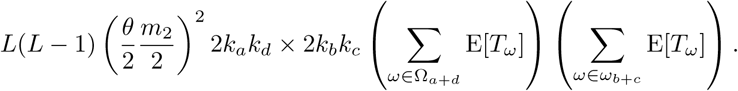

When the appropriate fourth order partial derivatives are taken we get

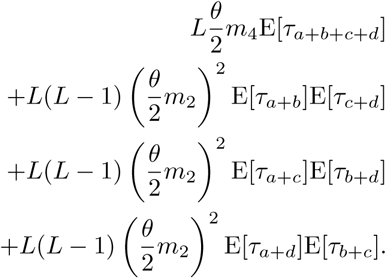

We can again simplify this expression for populations if we assume that individual are exchangeable. This gives

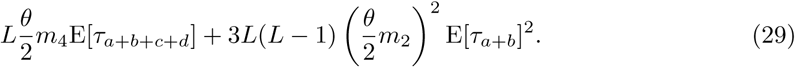

The expected kurtosis in the population is a quotient of random variables and calculating a second order approximation requires calculating eight order moments of the trait distribution. Instead we will calculate the expected fourth central moment.

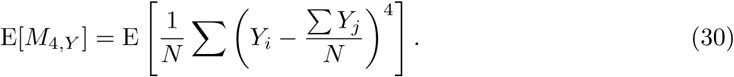

Examining the sum inside the expectation we see that

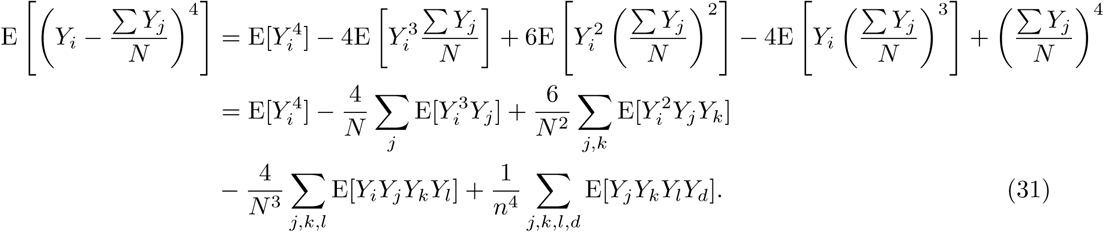

In calculating these expectations we have to remember that the value depends only on the number of times each variable appears in the expectation. That is, 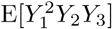 is equivalent to 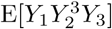 as long as all individuals in the population are exchangeable. The resulting expansion of (31) can be simplified by only considering terms of order one. Other terms can be ignored since we are assuming there are a large number of individuals in the population. This yields

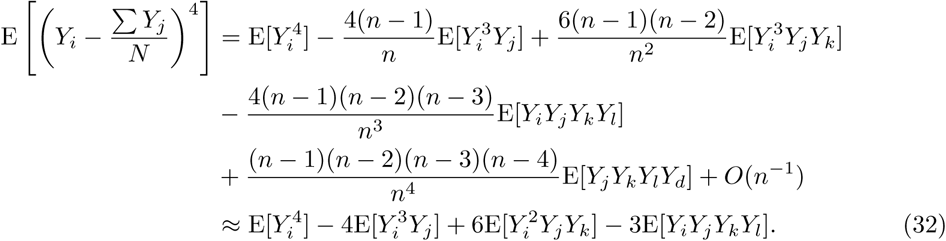

The first term, 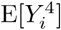 was derived in equation (25) as

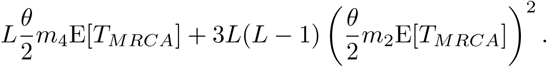

The second term, 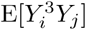 was derived in equation (26) as

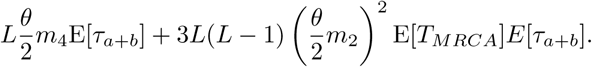

The third term, 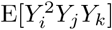 was derived in equation (27) as

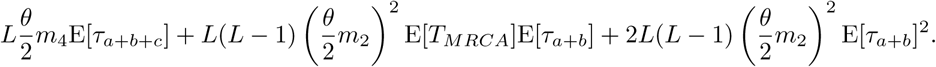

The fourth term, E[*Y*_*i*_*Y*_*j*_*Y*_*k*_*Y*_*l*_] was derived in equation (29) as

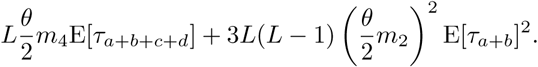

Plugging these into (32) we get

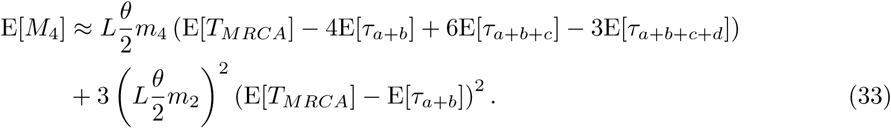

With some simple manipulation this can be rewritten in terms of 𝕋*_k,n_* to give

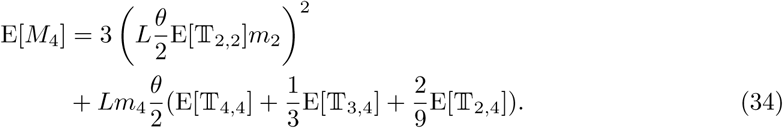

